# Biochemical and structural characterization of the RlaP family nucleotidyltransferase potentially involved in RNA repair

**DOI:** 10.64898/2025.12.29.696909

**Authors:** Amy Carruthers, Shirin Fatma, Raven H. Huang

## Abstract

Nucleotidyltransferases (NTases) of Polβ superfamily are highly diverse and play many important biological functions. However, many families of NTases remain uncharacterized. Because of physical proximity of the genes encoding NTases of PF10127 family to those encoding RNA ligases involved in RNA repair, PF10127 family of NTases was named RlaP (RNA ligase-associating Polβ). Here we report comprehensive characterization of two RlaP from *Pseudomonas fluorescens* and *Pseudomonas aeruginosa* (*Pf*RlaP and *Pa*RlaP), respectively. Our study showed that, among macromolecules isolated from *E. coli* cells, only RNAs are the substrates of RlaP. *In vitro* assays employing synthetic RNAs as substrates demonstrated that RlaP catalyzes addition of one or two nucleotide monophosphate (NMP) to the 3’-hydroxyl group of RNAs, with the damaged RNAs as the preferred substrates. The crystal structure of *Pf*RlaP provided insight into molecular recognition of RNA substrate and nucleotide triphosphate (NTP) by RlaP. Aminoacylation assays indicate that RlaP is required to restore biological function of the repaired tRNAs that have suffered excessive damage. Based on our studies, we propose an *in vivo* scenario where cell survival requires the involvement of RlaP in RNA repair to restore the biological function of the damaged RNAs.

## Introduction

The Polβ NTase superfamily is comprised of large and highly diverse families of proteins. Known NTases this superfamily transfer NMP or dNMP from NTP or dNTP to an acceptor hydroxyl group of a macromolecule or a small molecule. The members of the superfamily share the structural motif of NTase that possesses several conserved residues. Specifically, three conserved aspartate/glutamate residues coordinate two divalent ions for NTP or dNTP binding as well as for activating the acceptor hydroxyl group of a substrate for catalysis. Biologically, NTases play many important cellular functions, such as RNA polyadenylation by Poly(A) polymerases (*1*, *2*), tRNA maturation by CCA-adding enzymes (*3*, *4*), DNA repair by Polβ NTase (*5*), cell signaling by adenylate cyclase (*6*), innate immune signaling by cyclic GMP-AMP synthase (cGAS) (*7*), and bacterial CBASS signaling by CD-NTase (*8*).

Approximately twenty-five years ago, Aravind and Koonin performed the first major bioinformatic analysis of Polβ NTase superfamily (*9*), resulting in identification of three new NTase families. Fifteen years later, Kuchta *et al.* carried out a comprehensive classification of proteins possessing an NTase fold, with emphasis on identifying new human NTase families (*10*). One of the newly identified human NTases was later shown to be cGAS that produces cyclic GMP-AMP (cGAMP), an important signaling molecule in human innate immunity. More recently, during comprehensive bioinformatic analysis of proteins involved in RNA repair, Burroughs and Aravind have identified that genes encoding some NTases of the PF10127 family are in the same operons as the genes encoding a variety of RNA ligases (*11*). These NTases were therefore named RlaP (**R**NA **l**igase-**a**ssociating **P**olβ) and have been predicted to be involved in RNA repair. To date, there has been no report of the enzymatic activities or biological roles of the RlaP family NTases, which is comprised of more than 10,000 protein sequences.

Protein translation is essential for all organisms and is highly conserved. It is therefore a major target of biological toxins. Among cell-killing toxins targeting protein translation, ribotoxins that damage essential RNAs required for protein translation represent the largest group. To counter cell-killing by ribotoxins, some organisms and viruses employ RNA repair. This is exemplified by the action of the first RNA repair system from bacteriophage T4 discovered more than thirty years ago (*12*). We have also *in vitro* reconstituted the activity of Pnkp/Hen1 RNA repair system, which is widely distributed in bacteria (*13*). RtcB, a new class of RNA ligase, was discovered as the RNA ligase component for tRNA splicing in archaeal and eukaryotic organisms (*14*, *15*). The majority of RtcB, however, are found in bacteria. Because tRNAs in bacteria have no introns, it is reasonable to assume that most bacterial RtcB are likely to be involved in RNA repair (*16*, *17*).

Most cell-killing ribotoxins, such as PrrC (*12*), sarcin (*18*), γ-toxin (*19*), colicin E3 (*20*), colicin E5 (*21*), and colicin D (*22*), make a single cut of an essential RNA involved in protein translation to inhibit protein synthesis *in vivo*. To counter RNA damage by such “ordinary” ribotoxins, one would expect that direct RNA repair would be sufficient to restore the biological functions of the damaged RNA. Indeed, the T4 RNA repair system was able to restore *E. coli* tRNA^Lys^ cleaved by ribotoxin PrrC, preventing the T4-infected *E. coli* cells from committing suicide (*12*). In addition to these ordinary ribotoxins, however, “smart” ribotoxins have been identified. A “smart” ribotoxin is defined here as a ribotoxin that not only makes a cut on its RNA substrate, but also carries out additional enzymatic activities that result in loss of nucleotide(s) in the cleaved RNA. Two known “smart” ribotoxins are RloC and PaOrf2 (*23*, *24*). RloC first makes a cut on tRNA similar to ribotoxin PrrC, then excises the 3’-terminal nucleotide of the 5’-half of the cleaved tRNA (*23*). PaOrf2 cleaves tRNA like γ-toxin, followed by a second cut to remove two 3’-terminal nucleotides of the 5’-half of the cleaved tRNA (*24*). Therefore, due to the loss of nucleotide(s) on the damaged RNA by “smart” ribotoxins, it is uncertain whether such damaged RNAs can be repaired and still maintain their biological activities.

To provide insight into the enzymatic activity and biological function of RlaP, we cloned, overexpressed, and purified two RlaP from *Pseudomonas* organisms and performed *in vitro* reconstitution and structural studies. Our studies revealed that RlaP transfers one or two NMP to the 3’-hydroxyl group of RNA, and the enzyme possesses a certain degree of substrate specificity that is consistent with its involvement in RNA repair. In addition, we performed aminoacylation assays of the repaired tRNA to assess the requirement of RlaP for RNA repair. The collective studies described here led us to propose that RlaP could be the antidote of “smart” ribotoxins.

## Results

### Bioinformatic analysis of proteins of PF10127 family

To provide insight into the distribution of RlaP in a broader context of PF10127 family, we performed bioinformatic analysis of protein sequences of PF10127 family via construction of its Sequence Similarity Network (SSN) (Fig. 1A, Supplementary Fig. S1) (*25*, *26*). SSN of the PF10127 family revealed that the overwhelming majority of RlaP are from bacteria (Fig. 1A, colored blue). However, a small number of RlaP are also found in eukaryotic and archaeal organisms (Fig. 1A, colored red and green). Interestingly, a significant number of RlaP are from viruses (Fig. 1A, colored cyan), including bacteriophage T4 where the first RNA repair system was discovered (*12*). At Ecutoff value of 1e-30, the nodes of RlaP are distributed into eight major clusters, several minor clusters, and approximately forty isolated ones (Fig. 1A). The majority of RlaP are present in cluster 1, which can further be divided into sub-clusters 1a, 1b, and 1c (Fig. 1A). Both RlaP under current study belong to cluster 1a (Fig. 1A, marked with orange arrows), and the RlaP from bacteriophage T4 (T4NrdC.11) is present in cluster 5 (Fig. 1A, marked with a magenta arrow).

**Fig. 1.**
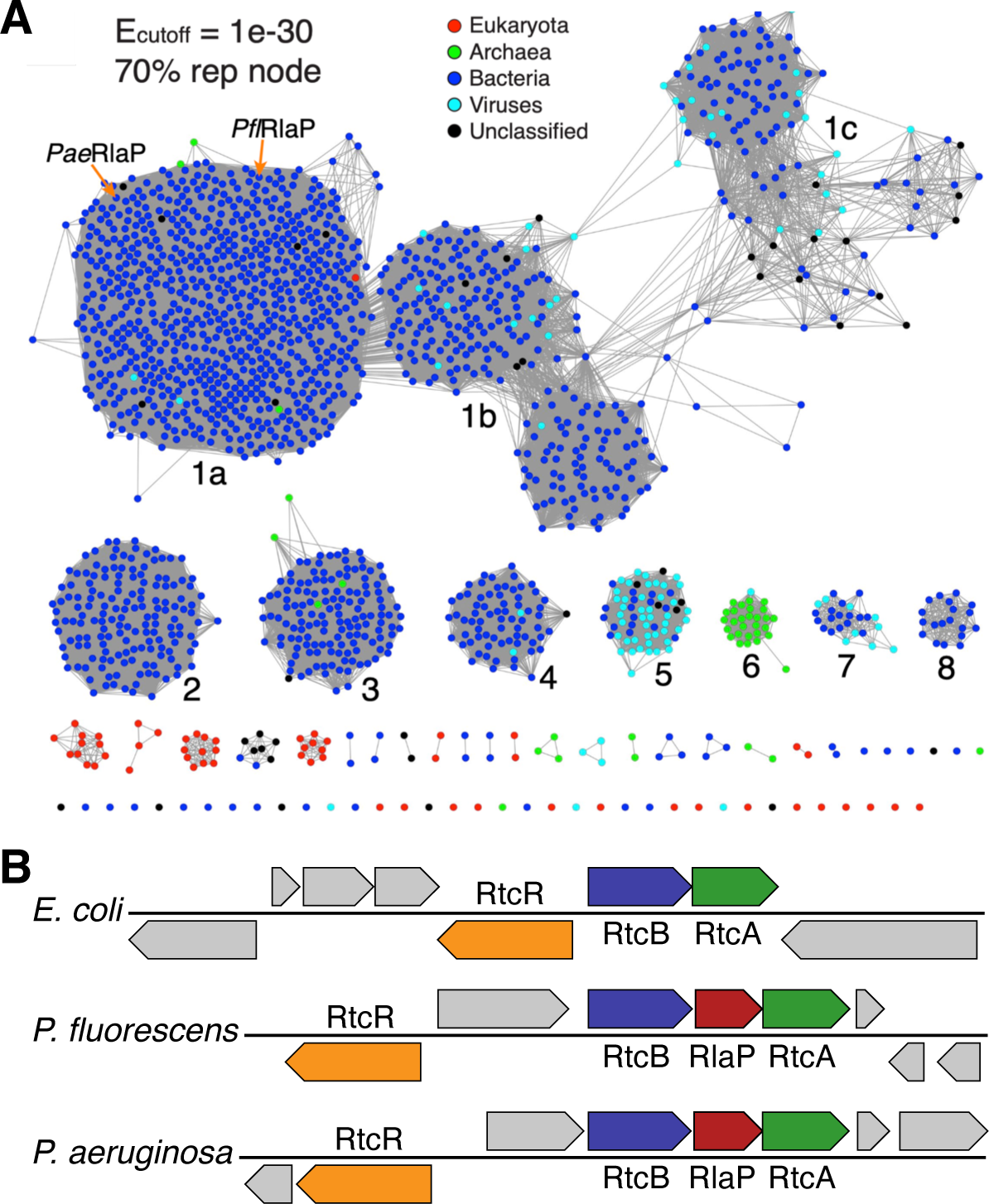
Distribution and gene neighborhoods of RlaP. (**A**) Sequence similarity network (SSN) of PF10127 family. The locations of two RlaP employed in this study are marked with orange arrows. And the location of RlaP of unknown function from bacteriophage T4 is marked with a magenta arrow. (**B**) Genome Neighborhood Diagrams (GND) of RtcB operons from three organisms. Only the genes involved in RNA repair as well as transcription activation are highlighted in color.

In *E. coli*, RtcB, a unique 3’-phosphate RNA ligases, is encoded in a two gene operon that also encodes RtcA (*27*), and operon is regulated by transcription activator RtcR (*27*, *28*) (Fig. 2B, top). In *P. fluorescens* and *P. aeruginosa*, similar gene organizations are also present (Fig. 2B, middle and bottom). The only exception is the presence of gene encoding RlaP, which is located between the genes encoding RtcB and RtcA (Fig. 2B, top). These two RlaP are the focus of current study. In addition to the frequent presence of the gene encoding RlaP in RtcB operons, it is also found in the operons of other RNA repair systems operons that encoding RNA repair systems the employ 5’-phosphate RNA ligases (Supplementary Fig. S2). *Pf*RlaP and *Pa*RlaP share 62% sequence identities (Supplementary Fig. S3). The recombinant *Pf*RlaP and *Pa*RlaP were overexpressed in *E. coli* and purified to homogeneity via several steps of chromatography (Supplementary Fig. S4).

**Fig. 2.**
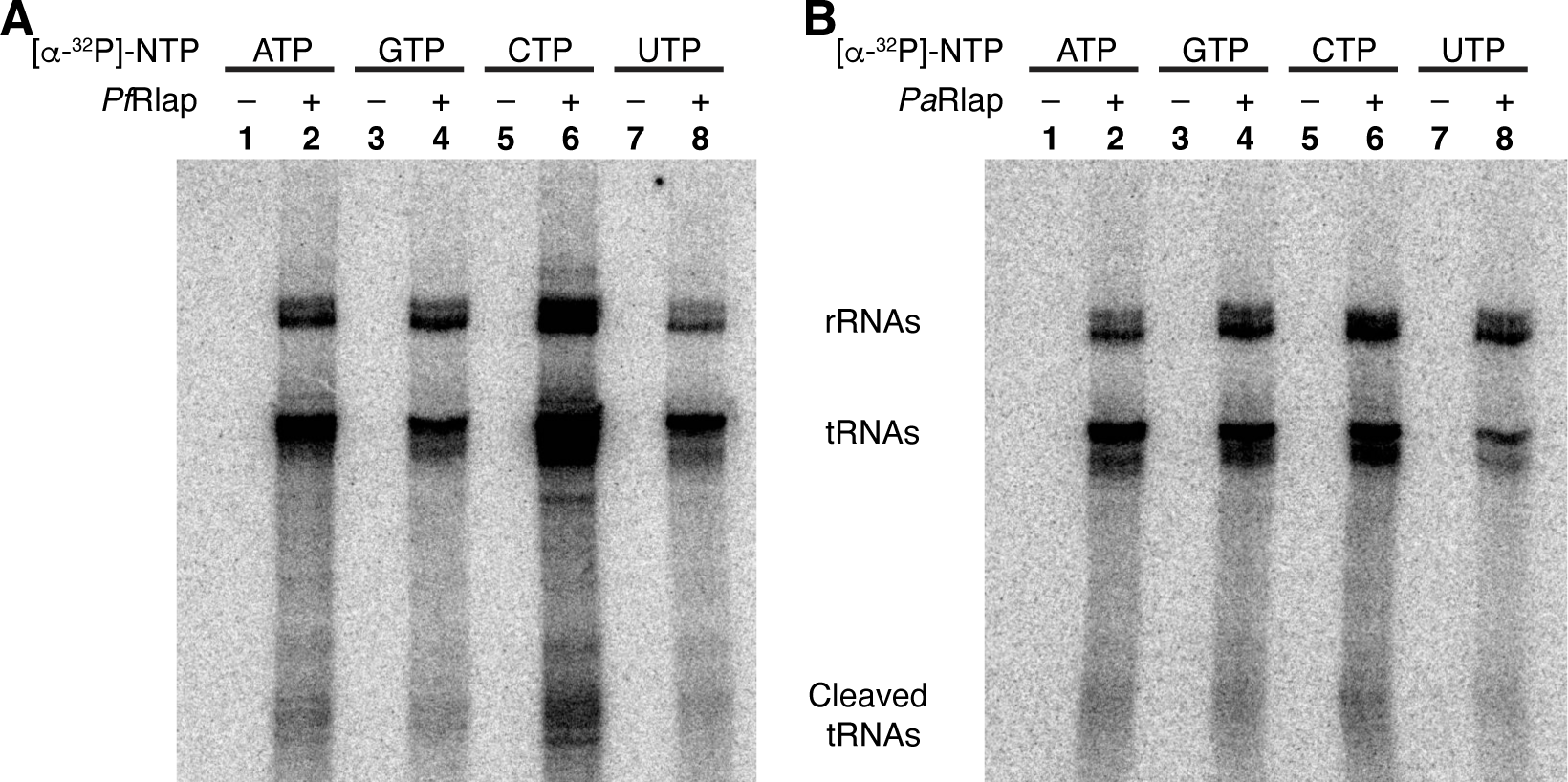
*E. coli* RNAs are the substrates of RlaP. *In vitro* reactions of total *E coli* RNAs and [α-^32^P]-NTP were carried out without and with the presence of *Pf*RlaP (**A**) and *Pa*RlaP (**B**).

### RNAs are the *in vivo* RlaP substrates

To shed light on potential *in vivo* RlaP substrates, we isolated total RNAs, genomic DNA, and proteins from *E. coli* cells and carried out reactions with [α-^32^P]-NTPs without and with the presence of *Pf*RlaP and *Pa*RlaP (Fig. 2, Supplementary Fig. S5 and Fig. S6). To compare the results with different substrates quantitatively, the reactions using total *E. coli* RNAs, DNA, and proteins were carried out under identical conditions, and three gels analyzing those reactions were exposed to a single imaging plate. This is to ensure that the images from three gels were at the same level of exposure, allowing the degree of reactions to be compared quantitatively.

As shown in Fig. 2, several *E. coli* RNAs matching the sizes of ribosomal RNAs, full-length tRNAs, and cleaved tRNAs were radiolabeled only in the presence of RlaP (Fig. 2, lanes 2, 4, 6, and 8), indicating that RNAs are the substrates of RlaP. Since all four NTPs can react with RNAs, we conclude that RlaP exhibits little substrate specificity for NTPs. We carried out the same reactions with *E. coli* genomic DNA and did not observe significant reaction (Supplementary Fig. S5), indicating that *E. coli* genomic DNA is unlikely a substrate of RlaP *in vivo*. Finally, we also carried out the reactions with *E. coli* proteins and concluded that *E. coli* proteins are also not the substrates of RlaP (Supplementary Fig. S6).

### *In vitro* reconstitution of RlaP activity using synthetic RNAs

To provide chemical and mechanistic insight into the reactions catalyzed by RlaP, we employed two synthetic RNAs, which mimic the product of tRNA_Asp_ cleavage by colicin E5 when annealed together, as potential substrates (Fig. 3A). In addition, each RNA has three variants depending on whether the terminal hydroxyl groups at the 5’- and 3’-ends are phosphorylated. They are unphosphorylated at both ends (34 and 42), 5’-phosphorylated (P34 and P42), and 3’-phosphorylated (34P and 42P). These six RNAs, either alone or annealed in various combinations to form mimics of the cleaved tRNA_Asp_, were subjected to the reactions carried out by *Pf*RlaP and *Pa*RlaP in the presence of [α-32P]-ATP (Fig. 3B).

**Fig. 3.**
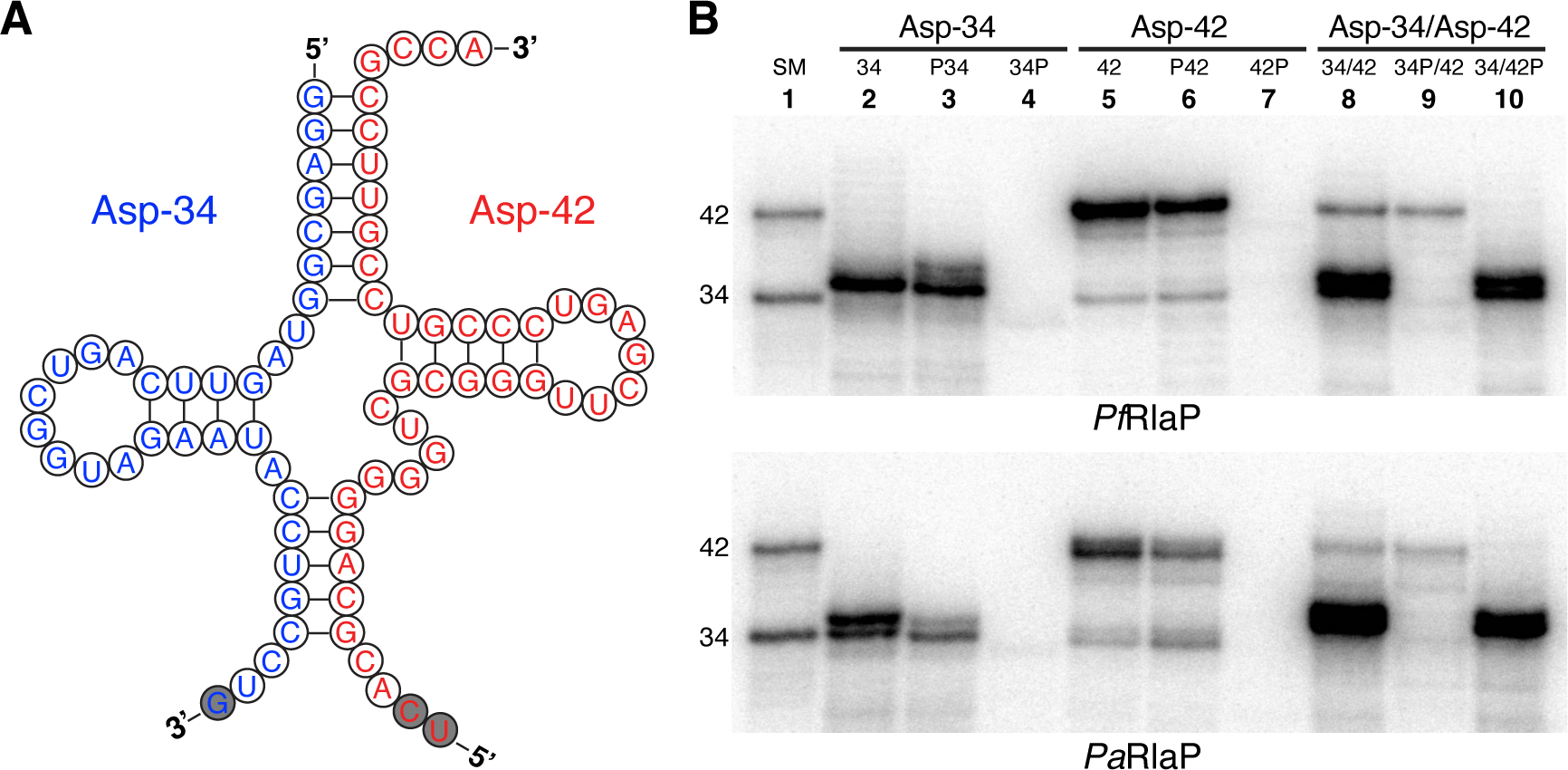
*In vitro* reconstitution of the enzymatic activity of RlaP using synthetic RNAs as substrates. (**A**) Sequences of two synthetic RNAs, which match the sequences of 5’- and 3’-halves of tRNA^Asp^ resulting from the cleavage by ribotoxin colicin E5. The nucleotides belonging to the anticodon of the tRNA are shaded gray. (**B**) DPAGE analyses of the reactions catalyzed by RlaP using various combinations of RNAs shown in **A** as substrates in the presence of [α-^32^P]-ATP. SM, size marker.

Single-stranded RNAs with either unphosphorylated or 5’-phosphorylated ends are effective substrates of RlaP (Fig. 3B, lanes 2, 3, 5, and 6). On the other hand, single-stranded RNAs with 3’-phosphorylated ends are not substrates of RlaP (Fig. 3B, lanes 4 and 7). These data indicate that effective RNA substrates of RlaP require the presence of 3’-OH group but not 5’-OH group. Similar results were observed with the reactions employing the annealed Asp-34/Asp-42 RNAs as substrates. Specifically, RlaP only catalyzes reactions on RNAs that possess 3’-OH groups (Fig. 3B, Asp-34 in lanes 8 and 10, and Asp-42 in lanes 8 and 9).

The requirement of 3’-OH group in RNAs to be effective substrates of RlaP indicates that, like other known families of NTases, RlaP likely catalyzes addition of 5’-NMP to the 3’-OH group of RNAs. This was further confirmed by the experiment of two-dimensional TLC analyzing nucleotides from the digested tRNAs (Supplementary Fig. S7). Judged by the mobilities of the products when compared to the size marker, RlaP adds one or two nucleotides to the RNA substrates (Fig. 3B). This was further confirmed by MALDI-MS experiments of the RNA products from the reactions catalyzed by *Pf*RlaP in a preparative scale (Supplementary Fig. S8). The mechanism of preventing RlaP from adding more than two NMP at the 3’-end of an RNA substrate (e.g., RlaP acts as an RNA polymerase) is unknown and requires further investigation.

Importantly, the relative reactivities of the same RNA substrate differ, depending on whether the RNA is alone or is part of a mimic of the cleaved tRNA. The reactivity of Asp-34 increases significantly from the single-stranded RNA to the mimic of a cleaved tRNA (Fig. 3B, compare lanes 2, 3 to lanes 8, 10). On the other hand, the opposite is true for Asp-42 (Fig. 3B, compare lanes 5, 6 to lanes 8, 9). These observations indicate that RlaP possesses a certain degree of RNA substrate specificity. Specifically, when presented with equal opportunity of reacting with substrate of 34/42, RlaP significantly prefers to catalyze the reaction on Asp-34 to Asp-42 (Fig. 3B, lane 8). This strongly suggests the involvement of RlaP in RNA repair *in vivo*, as the 3’-end of Asp-34 mimics one of the ends of the damaged tRNA generated by colicin E5.

### *In vitro* reactions using synthetic DNAs and dNTP

To further probe the substrate specificity of RlaP, we carried out the reactions catalyzed by RlaP using synthetic DNAs as potential substrates. The rationale for the experiments was that, although *E. coli* genomic DNA was not shown to be the substrate of RlaP (Supplementary Fig. S5), the negative result might stem from low availability of the 3’-OH end of DNA due to very large size of the genomic DNA. We thus purchased and purified two synthetic DNAs, D34 and D42, whose sequences are identical to the two synthetic RNAs we employed for our *in vitro* reconstitution (Fig. 3A). These two synthetic DNAs, either alone (D34 and D42) or annealed with each other (D34/D42) were subject to the reactions catalyzed by RlaP in the presence of [α-^32^P]-ATP (Fig. 4, top panel). In addition to DNA substrates, we also employed DNA/RNA hybrids (42/D34 and D42/34) for the reactions, and RNA substrates (34, 42, and 34/42) were used as the positive controls. Also, we tested the reactions with [α-^32^P]-dATP instead of [α-^32^P]-ATP (Fig. 4, bottom panel).

**Fig. 4.**
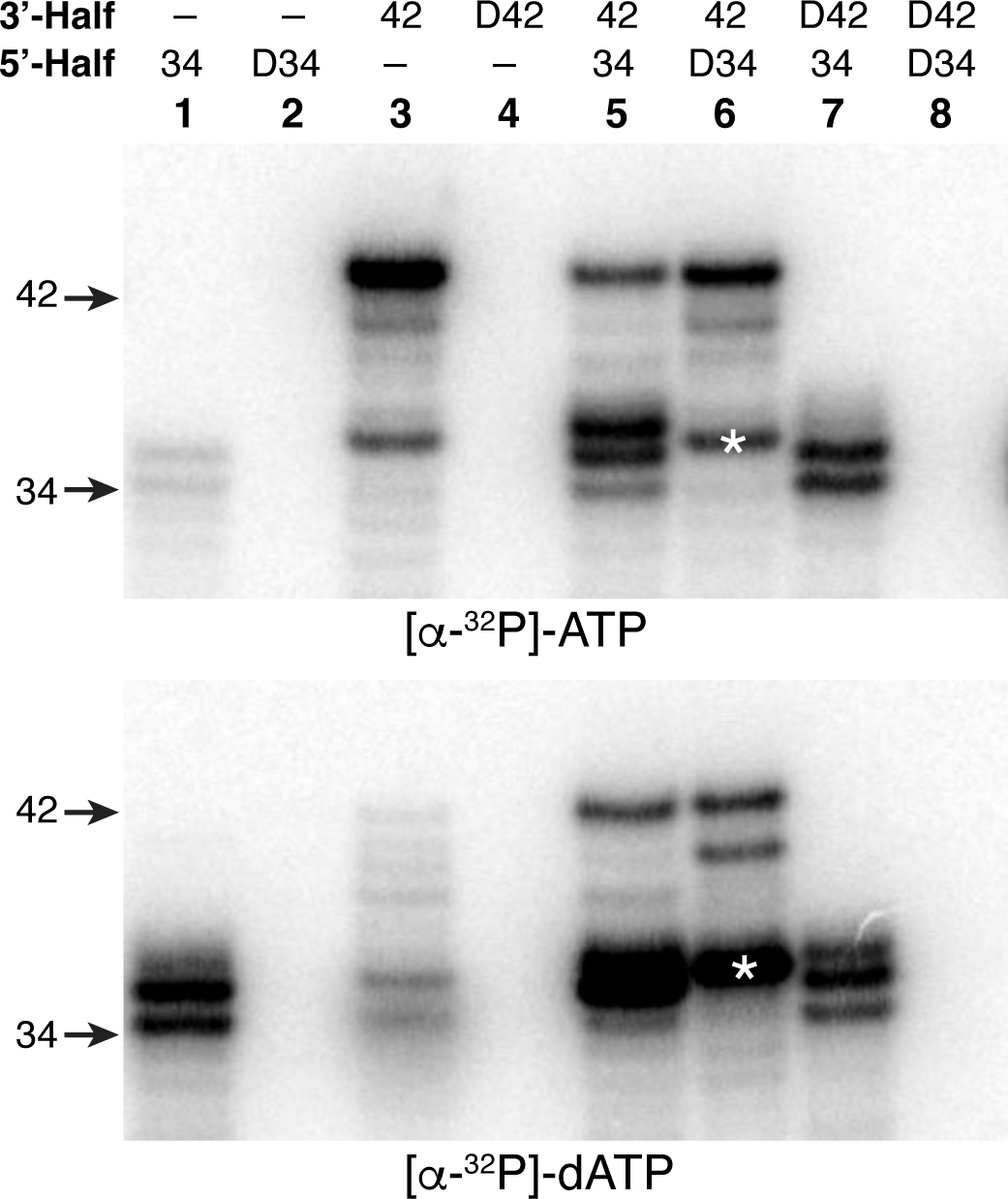
*In vitro* assays of the enzymatic activity of *Pf*RlaP using both synthetic DNAs and RNAs as substrates in the presence of [α-^32^P]-ATP (top panel) and [α-^32^P]-dATP (bottom panel). The bands marked with asterisk are likely the reaction product of degraded Asp-42, whose intensities differ significantly between the two panels.

As expected, all RNAs react with ATP (Fig. 4, lanes 1, 3, and 5 of the top panel). On the other hand, no reaction was observed with DNA, whether alone (Fig. 4, lanes 2 and 4), or the annealed with each other (Fig. 4, lane 8). When a DNA is hybridized to an RNA, only the RNA strand reacts with ATP (Fig. 4, lanes 6 and 7). We also carried out the same set of reactions in the presence of [α-^32^P]-UTP instead of [α-^32^P]-ATP and revealed that RlaP does not catalyze the reaction between synthetic DNAs and UTP (Supplementary Fig. S9). This is to confirm that the two minor products observed in the reaction between *E. coli* genomic DNA and [α-^32^P]-UTP (Supplementary Fig. S5, lane 8) are not from *E. coli* genomic DNA. The experiments employing synthetic DNAs described here, together with the reactions using *E. coli* genomic DNA (Supplementary Fig. S5), clearly demonstrate that DNA is not the substrate of RlaP.

Surprisingly, RlaP is able to catalyze the reaction between RNAs and dATP efficiently (Fig. 4, bottom panel). Although the overall pattern of the reactions between ATP and dATP is similar, there are some subtle differences in terms of substrate specificity. For example, dATP is more reactive than ATP with Asp-34 alone (Fig. 4, lane 1), but less reactive with Asp-42 alone (Fig. 4, lane 3). On the other hand, dATP is significantly more reactive than ATP with a degraded product of Asp-42 when Asp-42 is annealed to D34 (Fig. 4, lane 6, products marked with asterisk). Like ATP, RlaP does not catalyze reactions between synthetic DNAs and dATP.

### Optimization of *in vitro* reaction catalyzed by RlaP

At the early stage of our study, the *in vitro* reactions catalyzed by the recombinant RlaP were sluggish, which prevented us from obtaining high quality products for MALDI-MS. Specifically, we carried out our initial reactions employing Mg^2+^ because Mg^2+^ is the divalent ion preferred by most known NTases. To improve the efficiency of the reactions catalyzed by RlaP, we first screened various divalent ions and found that Mn^2+^ and Co^2+^ are significantly better than Mg^2+^ (Fig. 5A, compare lanes 2 and 3 to lane 1). Zn^2+^ is also a better divalent ion compared to Mg^2+^ (Fig. 5A, lane 6), but it is less effective than Mn^2+^ or Co^2+^, and the reaction is also less specific. Due to these observations, combined with the consideration that the physiological concentration of Mn^2+^ is significantly higher than Co^2+^ and Zn^2+^, we conclude that the *in vivo* activity of RlaP is likely to be Mn^2+^-dependent. We also carried out reactions at different pH and found that RlaP is most effective under weak basic condition (Fig. 5B).

**Fig. 5.**
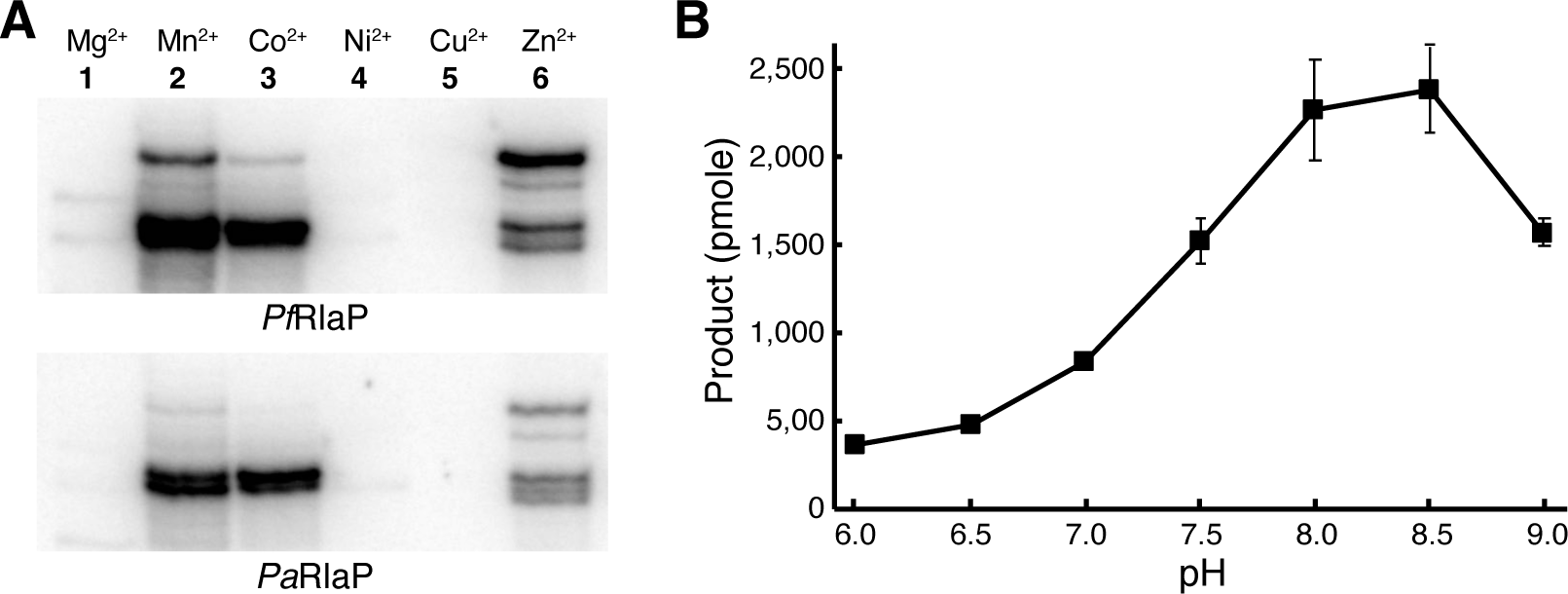
Optimization of the *in vitro* reactions carried out by RlaP. (**A**) Analysis of the reactions catalyzed by RlaP in the presence of different divalent ions. (**B**) The efficiency of the reactions catalyzed by *Pf*RlaP at different pH. The annealed 34/42 RNA was used as the substrates for the reactions.

### Crystal Structure of *Pf*RlaP

We crystallized and solved the structure of *Pf*RlaP at 2.1 Å resolution (Fig. 6, Supplementary Table S1). The overall fold of *Pf*RlaP can be depicted as the N-terminal NTase domain (Fig. 6A, colored red) fused with the C-terminal helical domain (Fig. 6A, colored blue). The interaction of these two domains is further reinforced by the helix at the very C-terminus of the enzyme (Fig. 6A, colored green), which joins the two helices from the N-terminal NTase domain to form a three-helix bundle. The overall fold of RlaP results in formation of a cleft between the N-terminal NTase domain and the C-terminal helical domain (Fig. 6C, D), where the RNA substrate and NTP presumably meet for the transfer reaction of NMP from NTP to the 3’-OH group of RNA. Although Mg^2+^ or Mn^2+^ were present in the crystallization solution, they were not observed in the final structure. Therefore, the NTP binding pocket shown in Fig. 6B is negatively charged, mainly reflecting the three strictly conserved aspartic acids required for catalysis. On the other hand, the bottom of the cleft, where the RNA substrate presumably binds, is mainly positively charged, (Fig. 6B, bottom). These assessments are consistent with the distribution of amino acid conservation of RlaP on the structure, which shows that the residues within the cleft are most conserved (Supplementary Fig. S10). In addition to the lack of divalent metal ions in the structure, attempts to include an NTP into the structure, either by co-crystallization or soaking, were also unsuccessful.

**Fig. 6.**
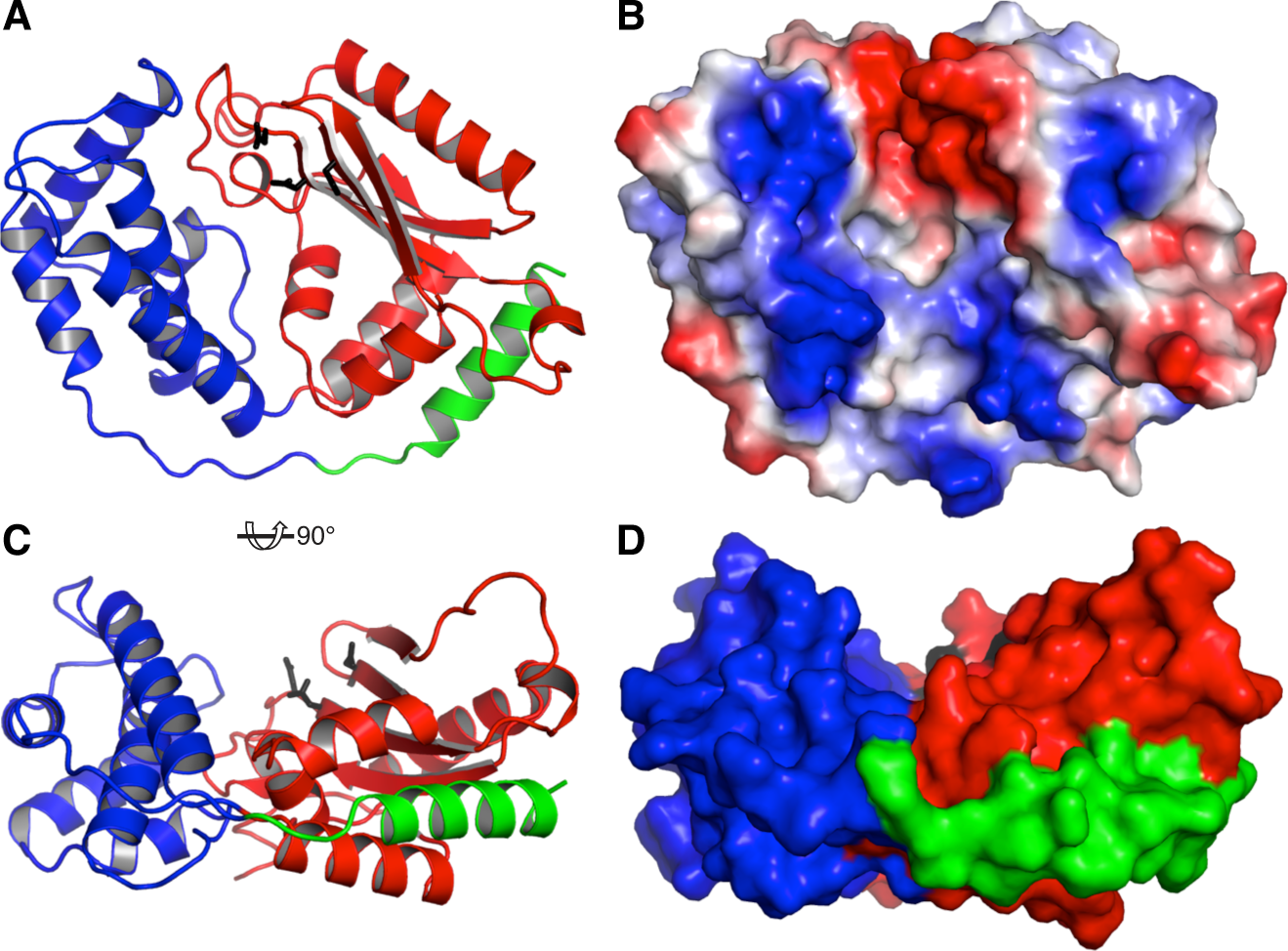
Structure of *Pf*RlaP-NTase. (**A**) Carton representation of the structure of *Pf*RlaP viewed from the top. The N-terminal NTase domain is colored red, the C-terminal helical domain is colored blue, and the very C-terminal helix joining the NTase domain is colored green. Three aspartic acids, required for the reaction and conserved in all NTase, are in stick and colored black. (**B**) Surface representation of the structure in the same view as in **A**. The structure is colored by surface charge, with negative charge in red, positive charge in blue, and neutral in white. (**C**) The side view of the same structure as in **A**. (**D**) The surface representation of the same structure as in **C**.

### Aminoacylation of the repaired tRNA^Asp^ by aspartyl-tRNA synthetase (AspRS)

As the first step to test the hypothesis that RlaP might be required for restoring biological activity of excessively damaged RNAs, we performed aminoacylation assays of various repaired tRNAs, differing in their treatments by RlaP prior to RNA repair (Fig. 7). Thus, two annealed tRNAs, Asp-34/P42 and Asp-33/P42 that mimic tRNA^Asp^ damaged by “ordinary” and “smart” ribotoxins, respectively, were treated with PfRlaP under various conditions (no NTP, GTP, and NTP). RNA repair was then carried out, and the repaired tRNA^Asp^ were purified for aminoacylation assays. The repaired tRNA^Asp^ from 34/P42, which should be the same as the wild-type tRNA^Asp^, was efficiently aminoacylated (Fig. 7, 34/P42 column). The efficiency of aminoacylation of the repaired 33/P42 is less than 10% of the repaired 34/P42 (Fig. 7, 33/P42 column), demonstrating that tRNA^Asp^ missing a nucleotide in the anticodon loop is a poor substrate of AspRS. Treatment of 33/P42 with RlaP in the presence of GTP prior to RNA repair results in significant increase of aminoacylation efficiency of the repaired tRNA^Asp^ (Fig. 7, 33/P42+GTP column), indicating that some of the repaired tRNA has been restored to the wild-type tRNA. Treatment of 33/P42 with RlaP in the presence of NTP mixture prior to RNA repair also increases the efficiency of aminoacylation by AspRS (Fig. 7, 33/P42+NTP column).

**Fig. 7.**
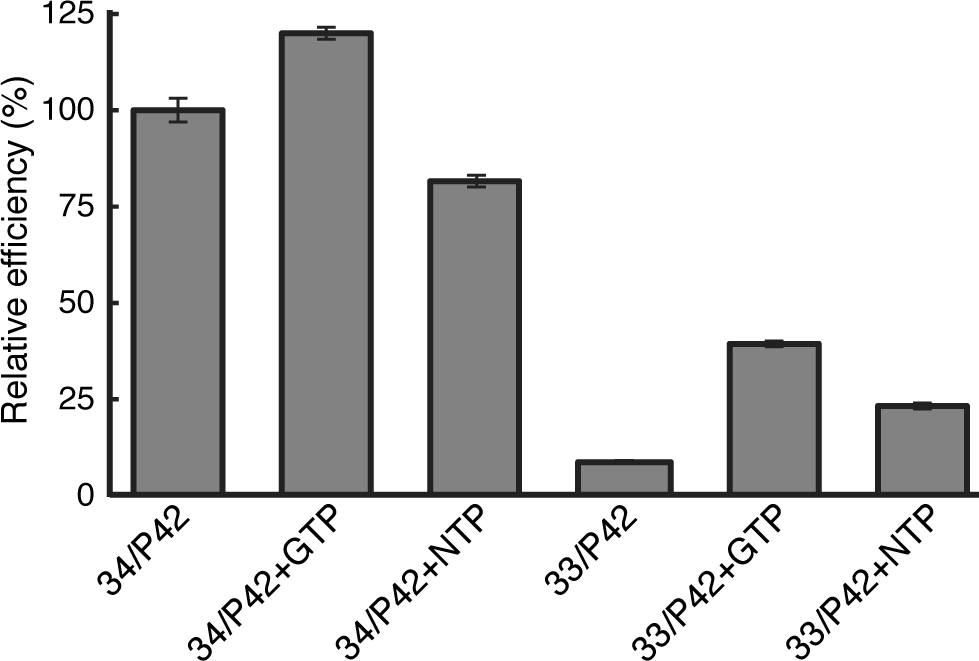
Efficiency of aminoacylation of various repaired tRNAs by AspRS. The repaired tRNA^Asp^ from 34/P42, which should be the same as the wild-type tRNA^Asp^, was used as the positive control, and the efficiency of its aminoacylation was normalized to 100%. All other assays were calculated as the efficiency relative to the one of the repaired 34/P42. The number of each assay shown is the average of triplicate experiments.

The aminoacylation assay alone, however, has an inherent flaw as a functional assay of RlaP. For example, the repaired tRNA resulted from the treatment of 34/P42 with RlaP in the presence of GTP showed approximately 20% increase in the aminoacylation efficiency by AspRS (Fig 7, 34/P43+GTP column), indicating that expanding the size of the anticodon loop of tRNA^Asp^ from the standard seven nucleotides does not significantly affect its aminoacylation. But the repaired tRNA is likely to be less effective in protein translation. Therefore, in addition to the aminoacylation assays described here, *in vitro* protein translation needs to be employed to test whether various aminoacyl tRNAs shown in Fig. 7 are effective substrates of protein translation. It is our judgement that the *in vitro* protein translation assays are too complicated and too technically challenging for us to be accomplished in a timely fashion for this publication.

## Discussion

Based on the data presented in this study, we propose mechanisms of cell death and survival via RNA damage and repair, using a tRNA as an example, as schematically depicted in Fig. 8. When a tRNA is damaged by an “ordinary” ribotoxin, a direct repair of the damaged tRNA is sufficient to produce biologically active repaired tRNA for cell survival (Fig. 8, pathway 1). When a tRNA is damaged by a “smart” ribotoxin such as RloC (*23*), however, there are two distinct possible outcomes of RNA repair. In the absence of a RlaP, a direct repair of a tRNA damaged by a “smart” ribotoxin would produce repaired, but biologically inactive tRNA, resulting in cell death (Fig. 8, pathway 2a). If RlaP is present in the cell, RlaP could react with the damaged tRNA, and an RNA repair system could concurrently carry out RNA repair. The combination of those two activities results in formation of heterogenous products of the repaired tRNA. The majority of the repaired tRNA is likely biologically inactive, as they either miss a nucleotide as shown in pathway 2a or the wrong nucleotide is incorporated by RlaP prior to RNA repair. However, a small portion of the repaired tRNA is expected to be biologically active due to the restoration of the correct missing nucleotide by RlaP prior to RNA repair (Figure 8, pathway 2b). We envision this could allow the cell to have a chance of survival in the emergency cases that require immediate action. The activity of RlaP would give the cell more time to initiate a more stringent cellular response to deal with the root cause of the crisis: RNA damage by a “smart” ribotoxin. Interestingly, reaction of RlaP with a damaged tRNA by an “ordinary” ribotoxin is likely to reduce the population of active tRNA after repair. We speculate that when the cell is in crisis mode with the invasion of a ribotoxin, its primary goal is to survive and the quantity of active tRNA is unlikely to be the main concern.

**Fig. 8.**
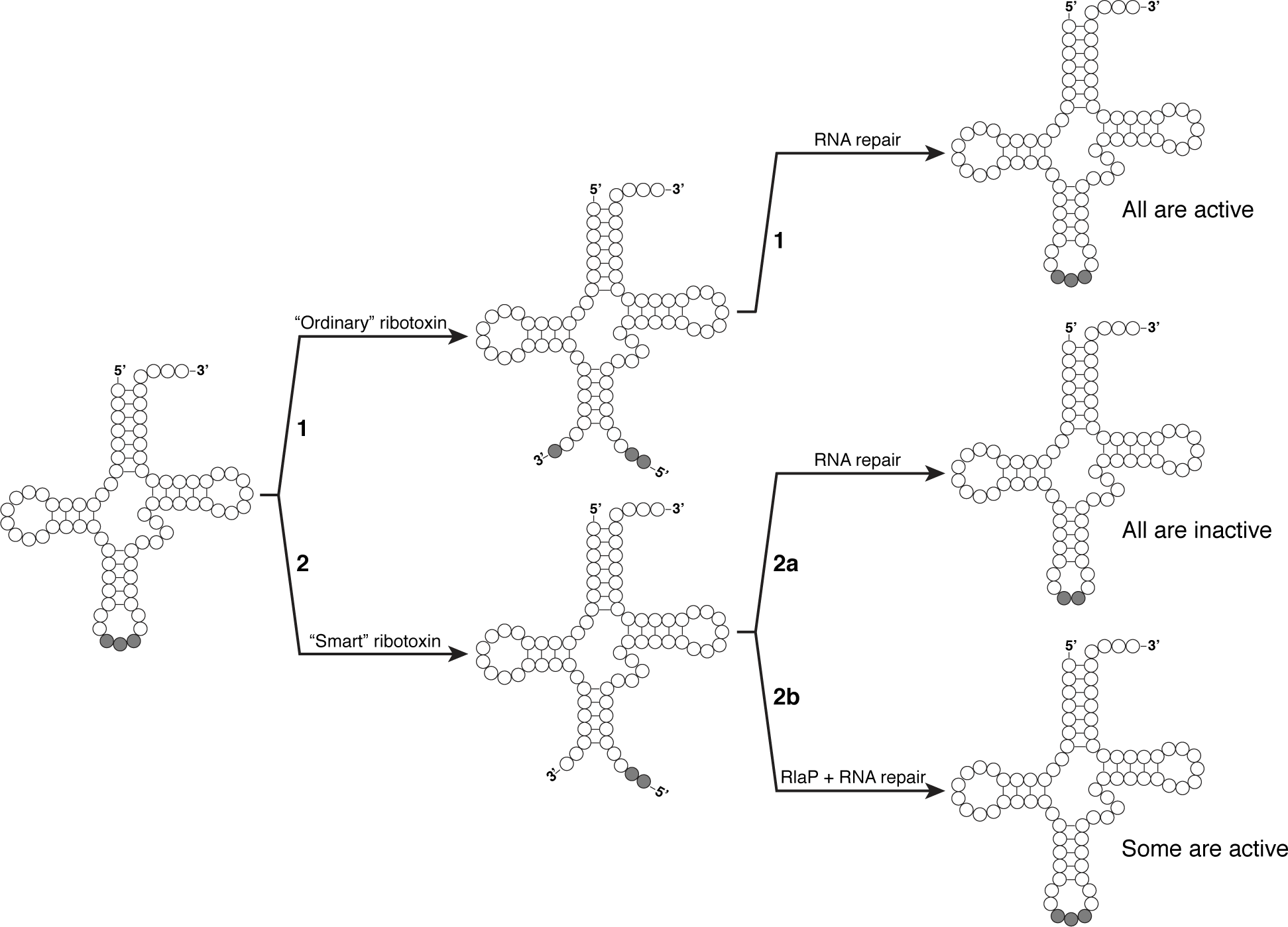
Proposed biological function of RlaP. Schematic view of a tRNA cleaved by ribotoxins, followed by RNA repair without or with the involvement of RlaP. The nucleotides corresponding to the anticodon of tRNA are highlighted in gray.

However, there are a couple of technical issues that need to be resolved for the proposed biological function of RlaP to be valid. Addition of one or two NMP to an RNA substrate results in the 3’-terminal nucleotide of RNA possessing a 3’-OH group. Repairing such an RNA is not an issue with the 5’-phosphate RNA ligases shown in Supplementary Fig. S2, as the reactions of those RNA ligases require an RNA with 3’-unphosphorylated end and a 5’-phosphorylated end. On the other hand, RtcB is a 3’-phosphate RNA ligase, requiring a 3’-phosphorylated end for RNA ligation. This in theory can be addressed by an RNA 3’-kinase, which is yet to be discovered. Alternatively, addition of two NMP to an RNA, followed by hydrolysis of the 3’-terminal nucleotide, should also produce the net addition of one NMP with the 3’-phosphorylated end, allowing RtcB to carry out RNA repair. Further study is required for the possible *in vivo* solution by bacteria of the incompatibility of RNA product of RlaP and its follow-up RNA repair by RtcB.

## Materials and Methods

### Expression and purification of RlaP

The RlaP genes from *pseudomonas fluorescens* and *pseudomonas aeruginosa* were purchased from IDT as gBlock Gene Fragments. The DNA fragments were digested by restriction enzymes and then inserted into pETDuet-1 vector using T4 DNA Ligase (New England Biolabs). The resulting plasmids were used to overexpress *Pf*RlaP or *Pa*RlaP in *E. coli* strain BL-21 (DE3) by the addition of 0.5 mM IPTG. RlaP was expressed for 16 hours at 18 °C. Cells were harvested by centrifugation. *Pf*RlaP pellets were resuspended in 20 mM Tris-HCl pH 8.0, 10 mM NaCl, 2% glycerol and 1 mM DTT and then lysed by three passes through a French press at 1500 psi. *Pf*RlaP was purified using DEAE ion exchange, Heparin affinity, and Mono Q ion exchange chromatography. *Pa*RlaP pellets were resuspended in 20 mM Tris-HCl pH 8.0, 200 mM NaCl, 2% glycerol and 1 mM DTT, lysed by French press, and purified using Heparin affinity and Mono Q ion exchange chromatography. Both proteins were stored at -80 °C in Storage buffer (10 mM HEPES pH 7.0, 400 mM NaCl and 2% glycerol). An additional size-exclusion chromatography step was carried out for *Pf*RlaP used for crystallization.

### Identification of *in vivo* substrate

Total RNA was purified from *E. coli* DH5α cells using TRIzol reagent (Thermo Fisher Scientific) according to the manufacturer’s instructions. Total RNA was precipitated in 75% cold ethanol, pelleted by centrifugation, and resuspended in Reconstitution buffer (50 mM Tris-HCl pH 8.0, 150 mM NaCl, 5 mM MnCl_2_). Genomic DNA was isolated from *E. coli* DH5α cells using DNeasy Blood and Tissue kit (Qiagen) according to the manufacturer’s protocol. Genomic DNA was eluted from the column using Reconstitution buffer. Purified genomic DNA was incubated with RNase I (Thermo Fisher Scientific) for 30 minutes at 37 °C according to the manufacturer’s instructions. To isolate *E. coli* protein, 300 mL DH5α cells (OD = 0.4) were pelleted and resuspended in 5 mL of Reconstitution buffer. Additionally, RNase I and DNase I (Sigma Aldrich) and 5 mM MgCl_2_ were added to degrade RNA and DNA. The cells were lysed by three passes through a French press at 1500 psi. The lysate was centrifuged for 45 min at 15,000 rpm to isolate the soluble protein fraction. The resulting supernatant was concentrated in a 3K Amicon Ultra centricon (Millipore) to remove digested nucleotides and small molecules.

Reactions were set up in Reconstitution buffer containing either 1 µg total RNA, 0.5 µg genomic DNA or 1 µg total protein, 10 µM NTP spiked with 125 nM [α-^32^P]-NTP and either 5 µM *Pf*RlaP, *Pa*RlaP or Storage buffer. Reactions were incubated at 25 °C for 30 minutes. The reactions were stopped by addition of DPAGE loading dye and heating at 95 °C for 3 min. RNA and DNA reactions were run on 10% DPAGE gels and protein reactions were run on 12% SDS gel. Gels were dried and exposed together to a phosphorimaging plate and radioactivity was detected using a PhosphorImager system.

### *In vitro* reconstitution of RlaP activity using synthetic RNA

For *in vitro* RlaP reactions, RNA oligomers first were annealed by heating for 3 min at 95 °C then cooled quickly for 10 min on an ice bath. Unless stated otherwise, RlaP reactions were carried out with 5 µM annealed RNA substrate, 2.5 µM *Pf*RlaP or *Pa*RlaP, and 5 µM NTP spiked with 62.5 nM [α-^32^P]-NTP in Reconstitution buffer. Reactions were carried out for 30 min at 25 °C. DPAGE loading buffer was added and 2 µL of each 10 µL reaction was run on a 15% DPAGE gel. The gel was dried and exposed to a phosphorimaging plate and radioactivity was detected using a PhosphorImager system.

Reactions with DNA and dATP were carried out in the same way as RNA and NTP reactions. The reactions for metal dependency were carried out as described above with the exception that 5 mM MnCl_2_ was replaced by the metal indicated in Figure 5A.

### Preparation of repaired tRNAs for aminoacylation

Asp-P42 RNA was annealed to either Asp-34 RNA or Asp-33 RNA as described previously. Reactions in preparative scale (300 µL total volume) were carried out in Reconstitution buffer containing 20 µM Asp-34/P42 or Asp-33/P42, 10 µM *Pf*RlaP, and 400 units of RNase inhibitor. Each of the RNA mixture was reacted at three different conditions: no NTP, at the presence of 100 µM GTP, or at the presence of NTP mixture (25 µM each of ATP, GTP, UTP and CTP). After 1 hour of incubation at 25 °C, 20 µM ATP and 350 units of T4 RNA ligase 1 were added directly to the reaction solution to ligate two RNA fragments into the full-length tRNA. The ligation reaction was carried out for 1 hour at 25 °C, and the ligated tRNA product was purified by 15% DPAGE.

### Aminoacylation Assays

Prior to aminoacylation reaction, each tRNA was folded by incubating at 75 °C for 3 min, followed by addition of 2 mM MgCl_2_, once the solution was cooled to 65 °C. The folding sample was finally cooled to the room temperature. Aspartylation of the repaired tRNAs (10 µM) by AspRS was carried out in 40 mM HEPES, pH 7.5, 4 mM ATP, 8 mM MgCl_2_, 200 µM unlabeled aspartic acid, 50 µCi ^3^H-[2,3]-L-aspartic acid, and 1 µM of AspRS. After 15 min of reaction at 37 °C, tRNAs were recovered via TCA precipitation on filter papers pre-soaked with 10% TCA. The filter papers were washed 3 times with 10% TCA to remove unreacted aspartic acid, followed by one time wash with 100% ethanol. After air dry, the filter papers were subject to scintillation counting. The same experiment using the *in vitro* transcribed tRNA^His^ as the background control. All assays were repeated three times and the data from the sets of experiments were used for the preparation of Fig. 7.

### Protein crystallization, data collection and structure determination

*Pf*RlaP was crystallized using hanging drop vaporization by mixing *Pf*RlaP (5 mg/mL) with the well solution (in 1:1 ratio) containing 1.2-1.6 M sodium malonate, pH 6.5-7.5, at 18 °C. Crystals were briefly soaked in cryoprotection solution containing 1.5 M sodium malonate and 20% glycerol followed by the same solution with 40% glycerol. The cryo-protected crystals were then mounted and flash-frozen in liquid nitrogen. Native and single-wavelength anomalous dispersion (SAD) data were collected at the 21-ID beamlines at the Advanced Proton Source (APS). Data were processed using the HKL2000 program (*29*). Phase was determined based on selenomethionine single-wavelength anomalous diffraction data using the Phenix program (*30*). A partial model was automatically built using Phenix. The remaining model was manually built using the Coot program (*31*). Refinement was done using Phenix.

## Acknowledgments

We thank Z. Wawrzak for help with data collection, S. Nair for the expression plasmid of AspRS, and L. Aravind for helpful discussion on RlaP.

