## Supplementary Figures and Table for "Biochemical and structural characterization of the RlaP family nucleotidyltransferase potentially involved in RNA repair"

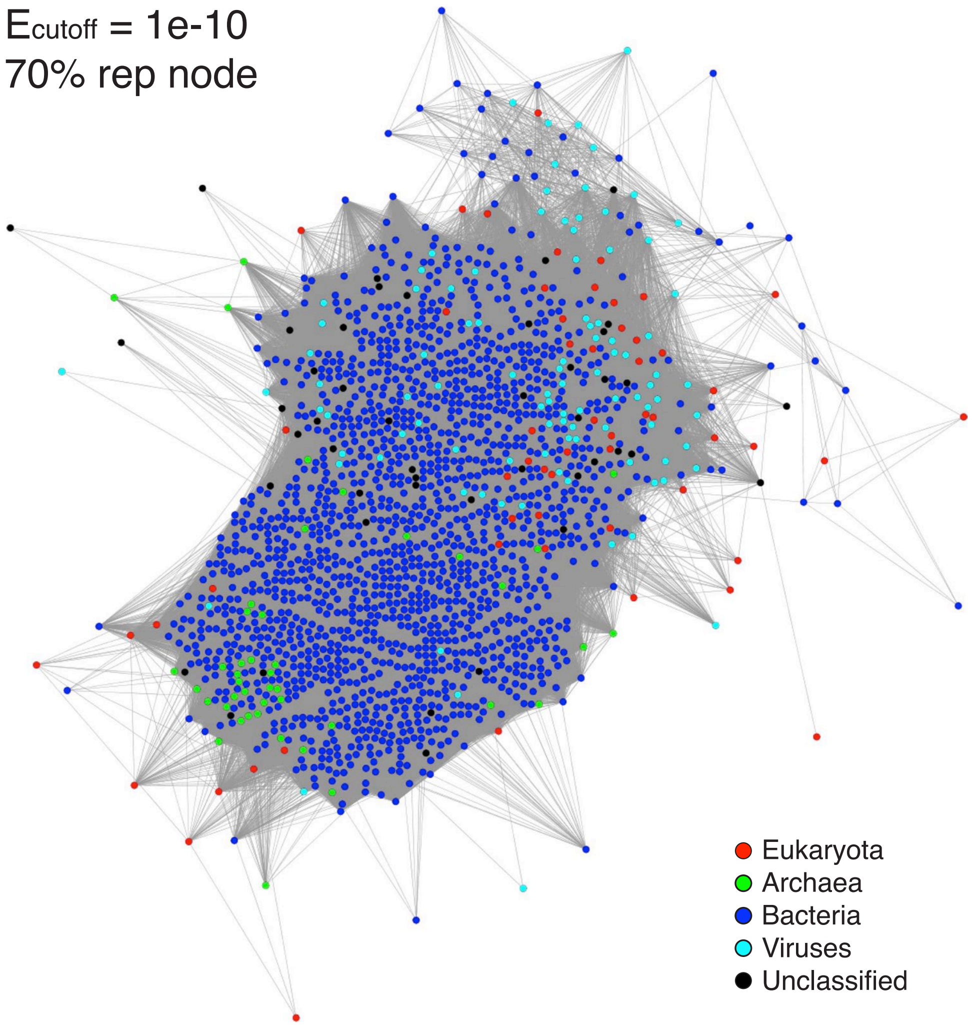

**Fig. S1. SSN of RlaP.** SSN is the same network as one shown in Fig. 1A except that it was calculated and displayed with $E_{cutoff}$ =1e-10. At $E_{cutoff}$ =1e-10, all nodes are included in a single cluster, indicating that all NTases of the PF10127 family are likely to carry out the same enzymatic reaction.

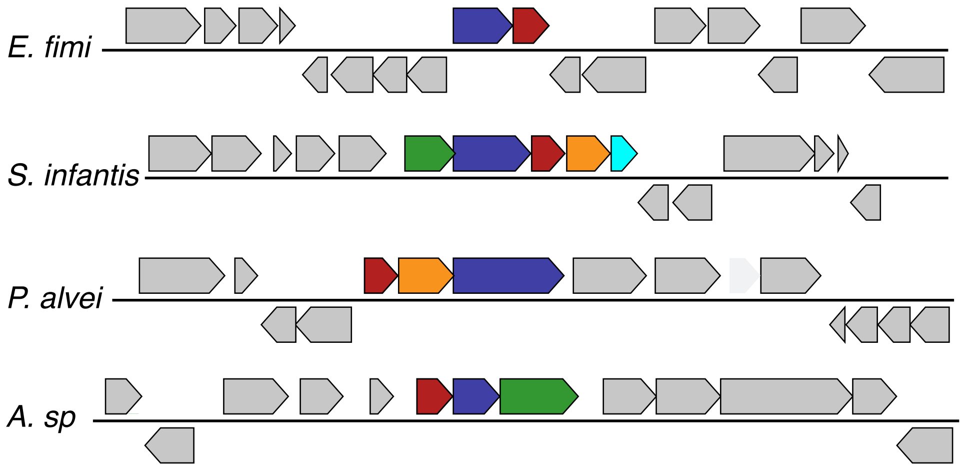

**Fig. S2. Examples of operons encoding 5’-phosphate RNA ligases and RlaP.** The operons encoding putative RNA repair systems are highlighted in color. RNA ligases and RlaP are colored the same as in Fig. 1B.

```
Score              Expect      Identities        Positives         Gaps
323 bits(828)      1e-116      166/266(62%)      199/266(74%)      4/266(1%)
PfRlaP 1    MQREERHPLSDAMRARVAQELDRIERERNVKVLYACESGSRAWGFASTDSDYDVRFVYVE  60
            M REERHPL  AMR RV  EL+RIERE +V VLYACESGSRAWGFAS DSDYDVRFVYV
PaRlaP 1    MNREERHPLPAAMRERVLDELERIEREHDVVVLYACESGSRAWGFASPDSDYDVRFVYVH  60

PfRlaP 61   KPDWFMQVDAPRDVIERPLDDELDVSGWELRKTLGLLRKSNPTLLEWLDSPLVYRQQTEA  120
            +P+W+ +V+ PRDVIERPL DELD+SGWELRK L L+RKSNP LLEWL SPLVYR++
PaRlaP 61   QPEWYQRVEEPRDVIERPLSDELDISGWELRKALRLMRKSNPALLEWLGSPLVYREEPGV  120

PfRlaP 121  TAQLRALAEAFYSPPAARNHYLSMARKNFRGYLQGETVRFKKYFYVLRPLLAVRWIDLGL  180
              +L  L  AF+S P +R+HYLSMARKN+RGYL+G++VR KKY YVLRPL AVRW+D GL
PaRlaP 121  REELLWLGSAFHSVPGSRHHYLSMARKNYRGYLKGDSVRLKKYLYVLRPLFAVRWLDAGL  180

PfRlaP 181  GRPPMTFADLL-STVSDAPLLDEVATLLALKRNAGEAAYGPRRPALHAFIVAEL---ERE  236
            G PP+ F  L+ +T+ D  L +E+  LL+LKR   E+AYGP R AL  FI AEL   ER
PaRlaP 181  GLPPVAFERLVEATLDDPSLREELDALLSLKRQRDESAYGPPRLALQRFIEAELLQAERG  240

PfRlaP 237  VPVLPRTREDSQRLDRYLRDTVKRFA  262
                   R+DS+ LDRYL D + ++
PaRlaP 241  ATQPTGARQDSRLLDRYLHDKIWQYT  266
```

**Fig. S3. Conservation between *Pf*RlaP and *Pa*RlaP.** The amino acid sequences of *Pf*RlaP and *Pa*RlaP were aligned, revealing 62% sequence identities between them. The conserved motifs of NTases, [GS], [D]h[D], and h[D]h (h represents hydrophobic residues), are in bold, and the residues within the conserved motifs required for catalysis are highlighted in red.

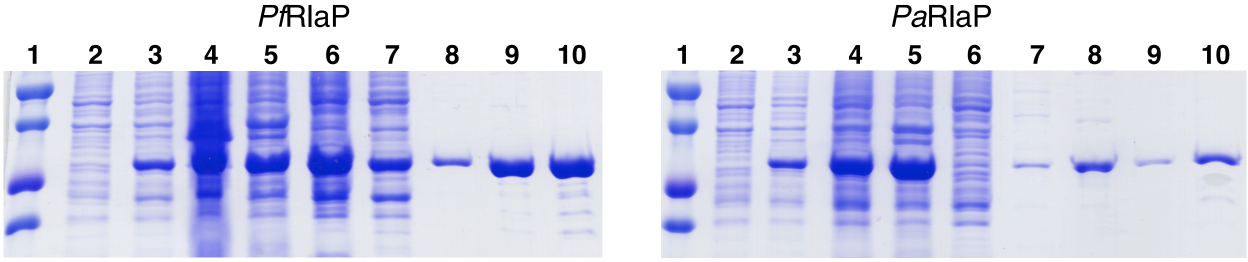

1. Size marker
2. Uninduced cells
3. Induced cells
4. Cell lysate
5. Cell pellet
6. Supernatant
7. Heparin
8. Mono Q
9. Gel filtration
10. Final concentrated protein

1. Size marker
2. Uninduced cells
3. Induced cells
4. Cell lysate
5. Cell pellet
6. Supernatant
7. Mono Q
8. Gel filtration loading
9. Gel filtration
10. Final concentrated protein

**Fig. S4. Expression and purification of the recombinant *Pf*RlaP and *Pa*RlaP.** Both proteins were expressed well in *E. coli* (lane 3). However, while more than half of the expressed *Pf*RlaP are soluble protein (lane 6 of the gel on the left panel), the overwhelming majority of the expressed *Pa*RlaP are insoluble (lane 5 of the gel on the right panel).

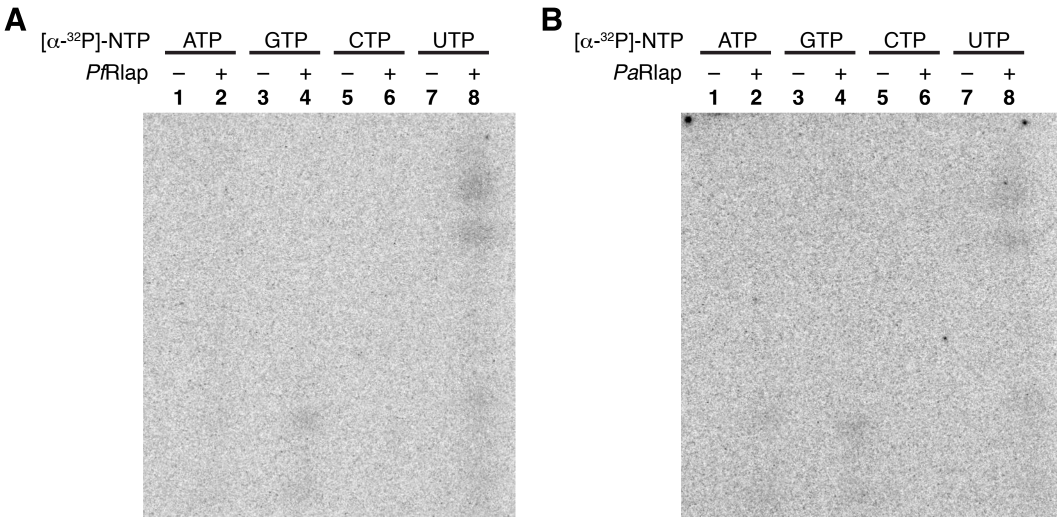

**Fig. S5. *E. coli* DNA is not the substrate of RlaP.** Genomic DNA isolated from *E. coli* cells was reacted with [α-$^{32}$P]-NTP in the absence (lanes 1, 3, 5, and 7) and presence (lanes 2, 4, 6, and 8) of *Pf*RlaP (**A**) and *Pa*RlaP (**B**).

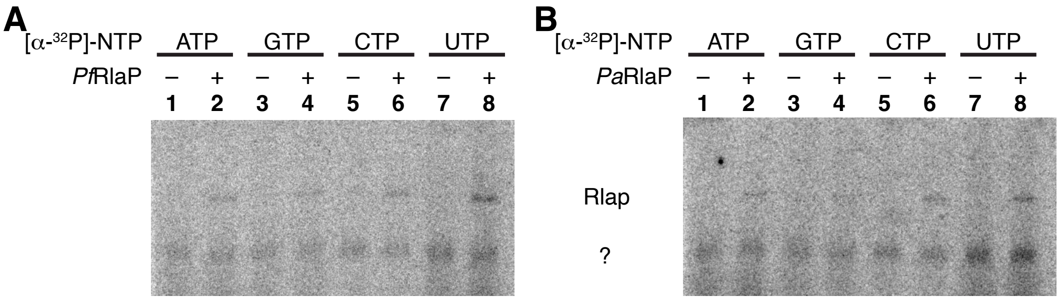

**Fig. S6.** ***E. coli*** **proteins are not the substrates of RlaP.** Proteins isolated from *E. coli* cells were reacted with [α-$^{32}$P]-NTP in the absence (lanes 1, 3, 5, and 7) and presence (lanes 2, 4, 6, and 8) of *Pf*RlaP (**A**) and *Pa*RlaP (**B**). Weak radioactive bands corresponding to the mobility of RlaP were observed in lanes 2, 4, 6, and 8, indicating possible formation of small amount of covalent complex of RlaP and NMP. The identity of RlaP, not an *E. coli* protein, was confirmed with the reactions in the absence of *E. coli* proteins (data not shown). In addition, an unknown molecule was radiolabeled (marked with a question mark). Because it is also present in the lanes lacking RlaP, we suggest that its formation is catalyzed by an *E. coli* enzyme.

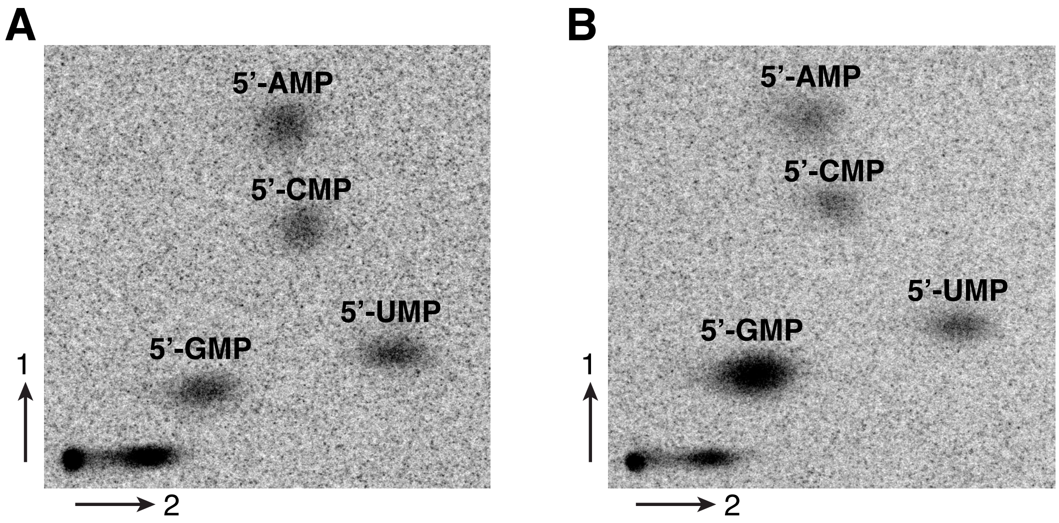

**Fig. S7. Confirmation of the addition of 5’-GMP to the 3’-OH group of RNA substrate by RlaP.** (**A**) 2D TLC of the digests of the *in vitro* transcribed $tRNA^{Gln}$. $tRNA^{Gln}$ was transcribed in the presence of [α-$^{33}$P]-NTP, resulting in radioactivity of all four nucleotides in tRNA. (**B**) 2D TLC of the digests of the same $tRNA^{Gln}$ as in **A**, but with the additional reaction with [α-$^{33}$P]-GTP catalyzed by *Pf*RlaP before the enzymatic digestion. Co-elution of 5-GMP from two sources of tRNA digests indicates that RlaP adds NMP to the 3’-OH group of the RNA substrate.

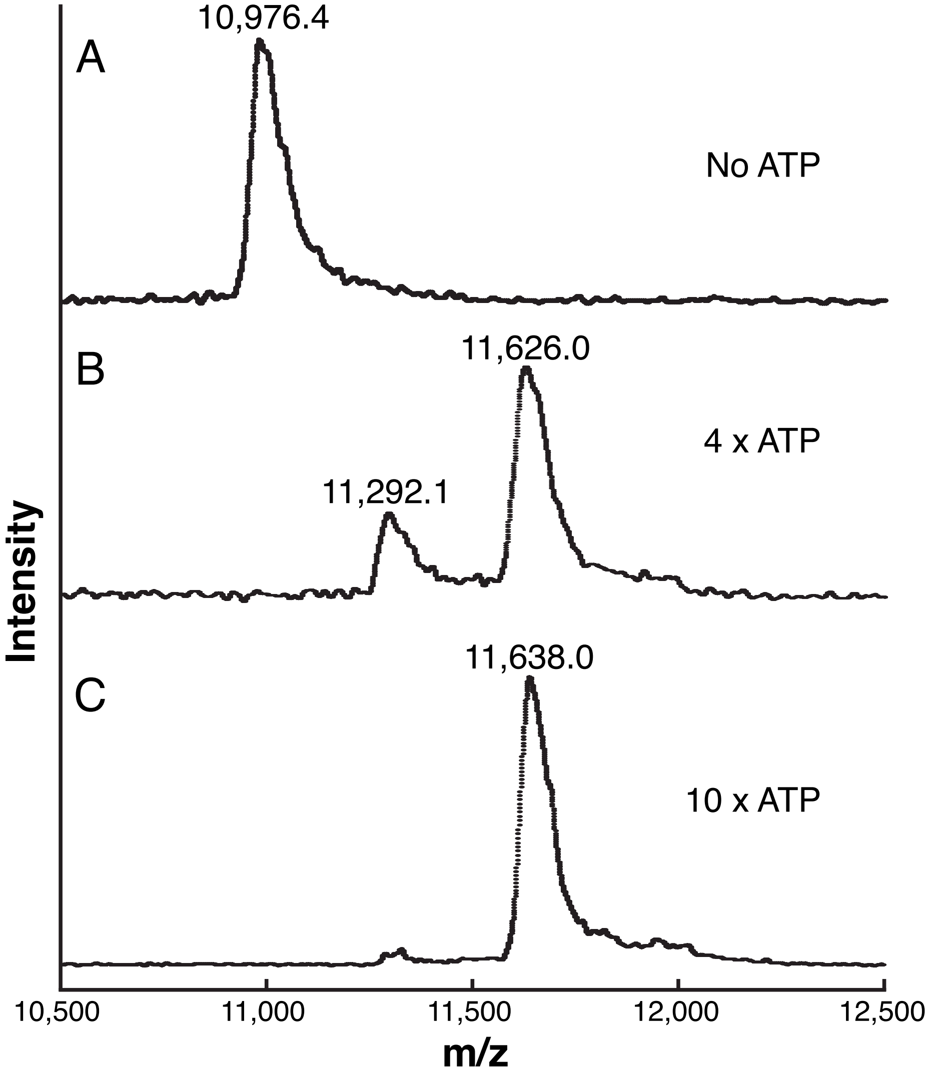

**Figure S8. MALDI-MS spectra of the reaction products of Asp-34 RNA.** 15 μM of Asp-34/Asp-42 RNA was incubated with 7.5 μM of *Pf*RlaP at 25 °C in the absence of ATP (**A**), in the presence of 60 μM ATP (**B**), and in the presence of 150 μM ATP (**C**). After 60 min of the reaction, RNA was purified with DPAGE. The recovered Asp-34 RNA was dissolved in water and subject to MALDI-MS experiments. The calculated m/z of Asp-34, Asp-34•AMP, and Asp-34•$(AMP)_2$ are 10,970.6, 11,317.0, and 11,665.0, respectively.

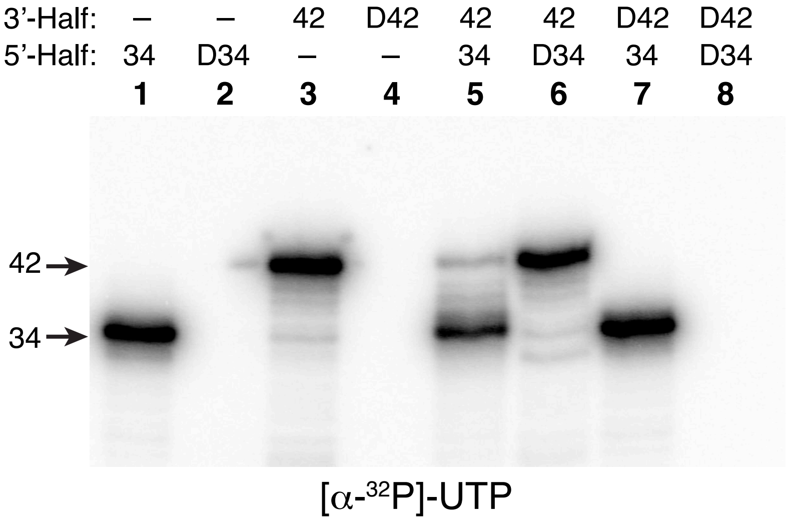

**Fig. S9. Assays of potential reactions of synthetic DNAs with [α-$^{32}$P]-UTP in the presence of *Pf*RlaP.** The reaction conditions are identical to those shown in Fig. 4, top panel, with the exception that [α-$^{32}$P]-UTP instead of [α-$^{32}$P]-ATP was used.

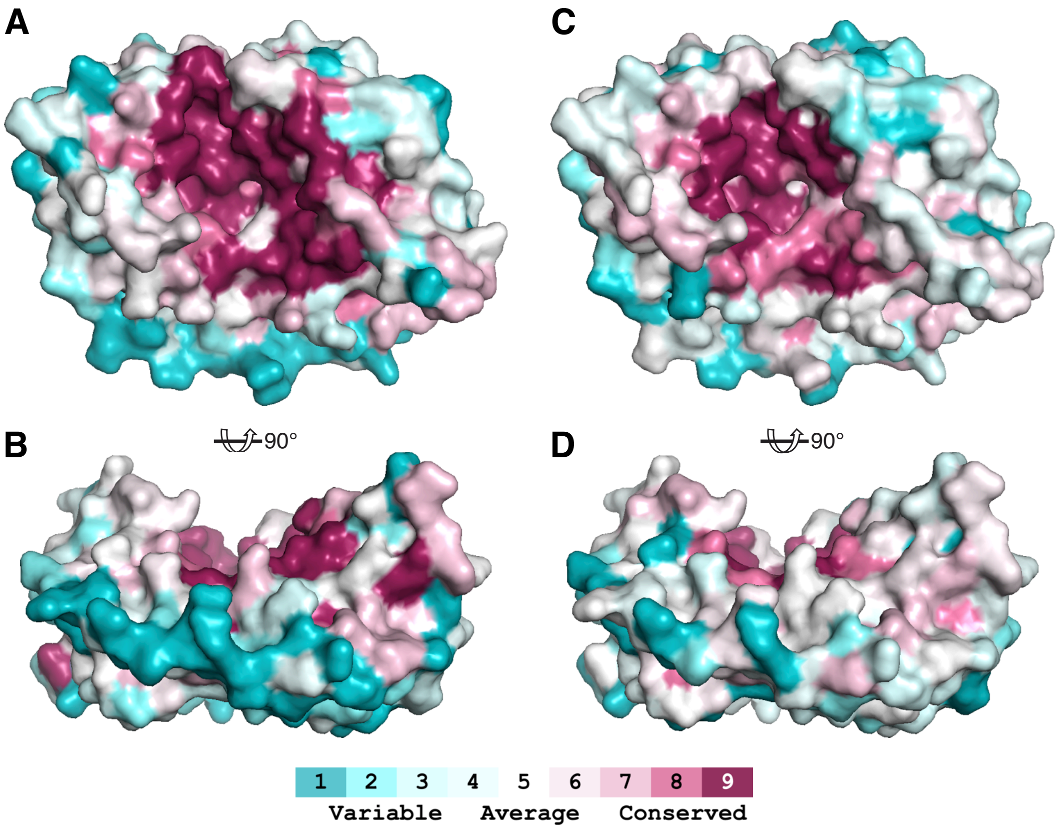

**Fig. S10. Amino acid conservation of RlaP of the PF10127 family depicted on the structure of *Pf*RlaP.** The structure of *Pf*RlaP was displayed and oriented the same as in Fig. 6B and Fig. 6D except the structure was colored based on amino acid conservation. The amino acid conservation in (**A**) and (**B**) was calculated based on the representative sequences from Cluster 1a of SSN shown in Fig. 1. On the other hand, the conservation shown in (**C**) and (**D**) represents the entire PF10127 family.

**Table S1. Summary of X-ray crystallography data collection and refinement**

| *Pf*RlaP | |
|---|---|
| **Data Collection** | |
| Space Group | $P2_12_12_1$ |
| Cell dimensions<br>a, b, c (Å)<br>α, β, γ (°) | <br>51.3, 69.7, 192.6<br>90.0, 90.0, 90.0 |
| Resolution (Å) | 50.0-2.1 (2.14-2.10) |
| $R_{merge}$ (%) | 6.4 |
| $I/\sigma$ | 11.1 (5.2) |
| Completeness (%) | 97.3 (99.7) |
| Redundancy | 5.7 |
| **Refinement** | |
| Resolution (Å) | 47.2-2.1 (2.18-2.1) |
| No. reflections | 40,128 (4,020) |
| $R_{work}/R_{free}$ (%) | 19.0/22.1 (19.4/22.7) |
| No. atoms<br>Protein<br>Ligand/ion | 4,614<br>4,272<br>342 |
| Average B factors (Å$^2$)<br>Protein<br>Water | <br>33.4<br>38.8 |
| R.m.s. deviations<br>Bond lengths (Å)<br>Bond angles (°) | <br>0.009<br>1.34 |
| Ramachandran statistics (%)<br>Favored<br>Allowed<br>Outliers | <br>98.1<br>1.9<br>0.0 |

*Values in parenthesis are for highest resolution shell.
